# Predictions from Deep Learning Propose Substantial Protein-Carbohydrate Interplay

**DOI:** 10.1101/2025.03.07.641884

**Authors:** Samuel W. Canner, Ronald L. Schnaar, Jeffrey J. Gray

## Abstract

It is a grand challenge to identify all the protein – carbohydrate interactions in an organism. Direct experiments would require extensive libraries of glycans to definitively distinguish binding from non-binding proteins. Computational screening of proteins for carbohydrate-binding provides an attractive and ultimately testable alternative. Recent computational techniques have focused primarily on which protein residues interact with carbohydrates or which carbohydrate species a protein binds to. Current estimates label 1.5 to 5% of proteins as carbohydrate-binding proteins; however, 50-70% of proteins are known to be glycosylated, suggesting a potential wealth of proteins that bind to carbohydrates. We therefore developed a novel dataset and neural network architecture, named **P**rotein **i**nteraction of **Ca**rbohydrates **P**redictor (PiCAP), to predict whether a protein non-covalently binds to a carbohydrate. We trained PiCAP on a dataset of known carbohydrate binders, and we selected proteins that we identified as likely *not* to bind carbohydrates, including DNA-binding transcription factors, cytoskeletal components, selected antibodies, and selected small-molecule-binding proteins. PiCAP achieves a 90% balanced accuracy on protein-level predictions of carbohydrate binding/non-binding. Using the same dataset, we developed a model named **Ca**rbohydrate **P**rotein **S**ite **I**denti**f**ier 2 (CAPSIF2) to predict protein residues that interact non-covalently with carbohydrates. CAPSIF2 achieves a Dice coefficient of 0.57 on residue-level predictions on our independent test dataset, outcompeting all previous models for this task. To demonstrate the biological applicability of PiCAP and CAPSIF2, we investigated cell surface proteins of human neural cells and further predicted the likelihood of three proteomes, notably *E. coli, M. musculus*, and *H. sapiens*, to bind to carbohydrates. PiCAP predicts that approximately 35-40% of proteins in these proteomes bind carbohydrates, indicating a substantial interplay of protein-carbohydrate interactions for cellular functionality.

**Significance Statement:** The totality of carbohydrate-protein interactions remains elusive, in part due to the inability to test proteomes versus glycomes in a high throughput manner. Here we show the first high-throughput methodology to predict protein-carbohydrate interactions at proteomic scales by using structural and sequence information. This information will allow scientists to target predicted protein-carbohydrate interactions to better determine how the elusive carbohydrate biomolecules play roles in all cellular functions.

## Introduction

In mammalian biology, carbohydrates are studied as two distinct families of molecules that are the focus of two disciplines. As metabolic precursors, from food or stored reserves, polysaccharides and monosaccharides (primarily glucose) are transported into and stored in the cytoplasm where they are subject to catabolic transformations to produce energy.^1^ In contrast, distinct covalent groupings of varied monosaccharide building blocks covalently bound to proteins (glycoproteins and proteoglycans) and lipids (glycolipids) are relatively stable and are abundant at the cell surface and in the extracellular milieu.^2^ A notable exception is O-GlcNAcylation, the reversible covalent attachment of the single sugar N-acetylglucosamine (GlcNAc) to serines and threonines of many cytoplasmic and nuclear proteins.^3^

Like their structures, the functions of carbohydrates are diverse. Among other functions they play essential roles in metabolism, they contribute to protein, cell and tissue structures, and they engage in molecular recognition upstream of cell-cell adhesion and cell regulation.^2^ Most of these functions involve engagement of glycans by proteins.^4^ These protein-carbohydrate interactions have predominantly been studied using chemical and biochemical methods, despite recent advances in the computational field.^5–7^

With the advent of the third generation of machine learning and large datasets, many novel algorithms have been created to better understand biophysical phenomena.^8,9^ Deep learning methods have recently overtaken most traditional algorithms for all biomolecular methods on all biopolymers, including prediction of protein structure, protein-small molecule interactions, and *de novo* protein design.^10–14^ Two of the largest computational steps in biophysics made recently are the releases of AlphaFold 2 (AF2)^10^ and ESM^15^. AF2 revolutionized the protein structure landscape by creating a public, easily accessible, and accurate method for protein structure prediction. AF2 additionally predicted the protein structures of 48 organisms that are publicly accessible.^12^ ESM (named for evolutionary scale modeling) revolutionized protein sequence representations through its transformer architecture, with ability to richly encode the language of protein sequences.^15^

Leveraging recent computational advances, scientists are beginning to explore the breadth of protein-carbohydrate interactions. We expect some of these protein-carbohydrate interactions to be involved in carbohydrate metabolism, some in intermolecular recognition and regulation of protein functions (e.g. O-GlcNAc), and others in cell adhesion and cell regulation. The goal of this work is to use computational advances to predict proteins amenable to carbohydrate binding, in its broadest interpretation. We use the term “lectome” to designate these proteins as a group, although here we identify proteins well beyond the original use of the term “lectin” on which it is based. The term lectin designates carbohydrate-binding proteins (other than antibodies), but excludes enzymes, carriers, or native sugar sensors. Here we computationally explore carbohydrate-binding proteins without excluding them based on function; we expect to capture proteins across metabolic, structural and molecular recognition functions. As discovery progresses, further sub-characterization may be of value.

Recently we developed a dataset and two models, named CArbohydrate Protein Site IdentiFier (CAPSIF):Graph and CAPSIF:Voxel, to predict the protein residues involved in noncovalent carbohydrate-protein interactions.^16^ CAPSIF:V and CAPSIF:G are trained and tested on the same datset and use the same residue level encodings, but CAPSIF:V encodes proteins onto a 3D voxelized grid with a UNet architecture whereas CAPSIF:G uses an equivariant graph neural network (EGNN) message passing framework; CAPSIF:V slightly outperformed CAPSIF:G by all measured metrics.

Since both CAPSIF models were released, two similar models have been created. Bibekar et al. released Protein Structure Transformer (PeSTo)-Carbs, which uses a geometric transformer architecture to predict residues involved in protein-carbohydrate interactions.^17,18^ PeSTo-Carbs employs a query-key-value attention mechanism with message passing across atoms that are then pooled for residue-wise predictions.^17^ He et al. released DeepGlycanSite, which leverages a geometric message-passing architecture to predict a glycan binding site in both the case of a known ligand and an unknown ligand.^18^ PeSTo-Carbs modestly outperforms both CAPSIF models on all reported metrics, whereas DeepGlycanSite focuses on binding to nucleotide structures as compared to carbohydrate-only polymers.

Most carbohydrate-protein interaction algorithms rely on multiple datasets to extract experimental coordinates for prediction.^16,17^ Currently the standard protein-carbohydrate dataset is UniLectin; however, UniLectin focuses only on proteins in the lectin family and thereby does not include other carbohydrate binding proteins.^19^ Recently, DIONYSUS was released detailing an immense set of experimental carbohydrate binding proteins with non-covalently bound carbohydrate as well as glycosylated proteins.^20^

Since experimentally solved structures can be difficult to obtain, especially in the presence of a carbohydrate ligand, some datasets of sequences exist that identify carbohydrate binding proteins. The Carbohydrate Active enZymes (CAZy) dataset identifies sequences of catalytically active proteins that act on glycosidic bonds.^21^ LectomeXplore is a dataset that identifies known lectins, their associated structures (if known) and potential lectins as identified by sequence similarity via a hidden Markov model (HMM).^22^ Rather than limiting their work to known lectins, Zhang et al. developed high throughput experiments with a ganglioside probe that identified 873 putative human proteins that likely interact with gangliosides.^23^ These works are limited by their scope, requiring either specific protein families or specific carbohydrate species to interact.

Here, we present novel frameworks to both predict whether a protein can bind to carbohydrates and where on that protein the carbohydrate binds, entitled **P**rotein **i**nteraction of **CA**rbohydrate **P**redictor (PiCAP) and **CA**rbohydrate **P**rotein **S**ite **I**denti**F**ier **2** (CAPSIF2). Both models leverage a large dataset with two training stages, first using all small molecule binding interfaces and then fine tuning with carbohydrate-specific data. We assess the ability of these models in their tasks. We then validate PiCAP against the work of Zhang et al.^23^ and identify potential outliers in their dataset. Finally, we use these models to make the first prediction of carbohydrate binding proteins, and residues of these proteins, of three proteomes. While these first proteome-wide predictions are likely noisy, they define the broad scope of the problem and invite refinement by future experimental and computational methods.

## Results

### NoCAP: a novel non-binder dataset

Many datasets exist for protein-carbohydrate interactions, with the most notable being DIONYSUS (Table 1). However, there is no dataset of proteins that do *not* bind to carbohydrates; therefore, we developed a novel dataset consisting of proteins known to bind carbohydrates and proteins that likely do not bind carbohydrates based on biophysical intuition. Although the non-binder dataset is likely mildly contaminated with some currently unknown carbohydrate binding proteins, we believe this dataset to be generally representative of proteins that do not bind carbohydrates. We denote this novel combined dataset as Nonbinder and binder of CArbohydrate Protein interactions (NoCAP) (Table 1). In addition, we created a subset of NoCAP, named DIONYSUS-Residue (DR) as all binding proteins in NoCAP with a bound ligand, retaining the DIONYSUS name as most protein structures were retrieved from the DIONYSUS dataset (Table 1).

**Table 1:**
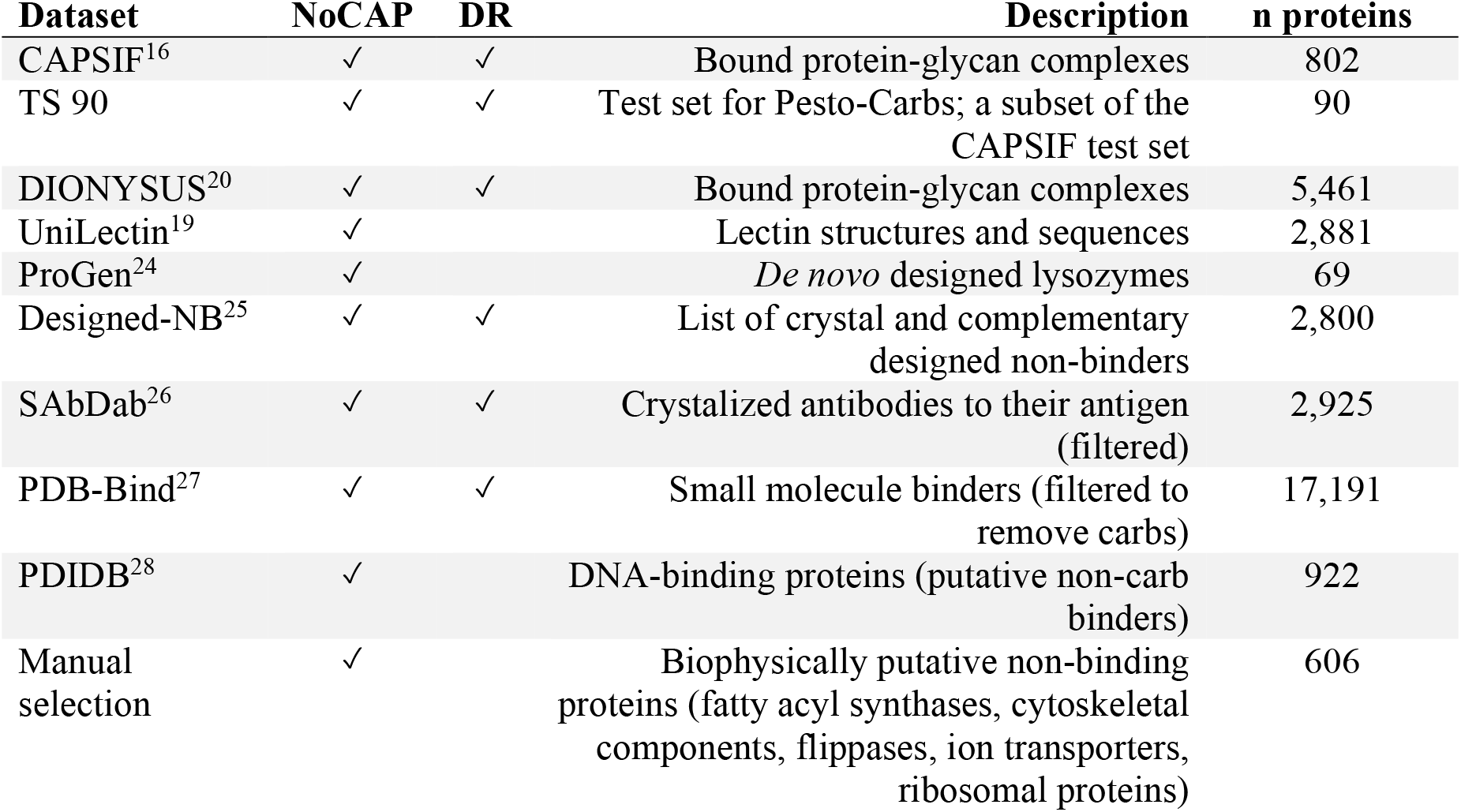
Experimental structural datasets. Columns 2 and 3 indicate dataset inclusion in our NoCAP or DR datasets.

| Dataset | NoCAP | DR | Description | n proteins |
| --- | --- | --- | --- | --- |
| CAPSIF<sup>16</sup> | ✓ | ✓ | Bound protein-glycan complexes | 802 |
| TS 90 | ✓ | ✓ | Test set for Pesto-Carbs; a subset of the CAPSIF test set | 90 |
| DIONYSUS<sup>20</sup> | ✓ | ✓ | Bound protein-glycan complexes | 5,461 |
| UniLectin<sup>19</sup> | ✓ |  | Lectin structures and sequences | 2,881 |
| ProGen<sup>24</sup> | ✓ |  | *De novo* designed lysozymes | 69 |
| Designed-NB<sup>25</sup> | ✓ | ✓ | List of crystal and complementary designed non-binders | 2,800 |
| SAbDab<sup>26</sup> | ✓ | ✓ | Crystalized antibodies to their antigen (filtered) | 2,925 |
| PDB-Bind<sup>27</sup> | ✓ | ✓ | Small molecule binders (filtered to remove carbs) | 17,191 |
| PDIDB<sup>28</sup> | ✓ |  | DNA-binding proteins (putative non-carb binders) | 922 |
| Manual selection | ✓ |  | Biophysically putative non-binding proteins (fatty acyl synthases, cytoskeletal components, flippases, ion transporters, ribosomal proteins) | 606 |

### CAPSIF2 outcompetes all previous models identifying carbohydrate-binding residues

We constructed an equivariant graph neural network (EGNN) named Carbohydrate Protein Site IdentiFier 2 (CAPSIF2) leveraging the same general architecture of our previous work CAPSIF:Graph (CAPSIF:G). Although CAPSIF:G underperformed CAPSIF:V, we chose the EGNN architecture because it is scalable to proteins of any size, while CAPSIF:V is limited by the size of the underlying convolutional voxels. Although the dataset of this work (6,724 protein structures) is substantially larger than our previous work (∼800 protein structures), there is still an intrinsic data imbalance in that most protein residues (∼95%) do not bind carbohydrates. To address this, we once again leveraged the Dice loss (Table 2) to emphasize the residues that bind carbohydrates (see methods).

**Table 2:** Average metrics for each deep learning architecture on test sets. Dice coefficient is described as 2TP / (2TP + FP + FN), where TP, FP, and FN are the counts of the true positives, false positives, and false negatives, respectively. MCC is the Matthews correlation coefficient. Boldface indicates the best performance for each metric.

| Model | DR Dice | DR MCC | TS 90 Dice | TS 90 MCC |
| --- | --- | --- | --- | --- |
| CAPSIF2 | **0.573** | **0.574** | 0.616 | 0.607 |
| Pesto-Carbs | 0.493 | 0.492 | **0.638** | **0.624** |
| CAPSIF:V | 0.226 | 0.202 | 0.608 | 0.622 |

In Figure 1 and Table 2, we compare our results to PesTo-Carbs^17^ and our previous model CAPSIF:V^16^. On the TS-90 test set, CAPSIF2 achieves 0.616 Dice and 0.607 MCC metrics and PesTo-Carbs outcompetes our model on this test set with a 0.638 Dice coefficient and 0.624 MCC (Table 2). Contrarily on the DR test set, CAPSIF2 achieves 0.573 Dice coefficient and 0.574 MCC, while PesTo-Carbs only achieves 0.493 Dice and 0.492 MCC metrics. On a per target basis, CAPSIF2 performs greater than 0.15 Dice better than PesTo-Carbs on 40% of targets and PesTo-Carbs performs greater than 0.15 Dice than CAPSIF2 on 15% of targets (Figure 1B).

**Figure 1:**
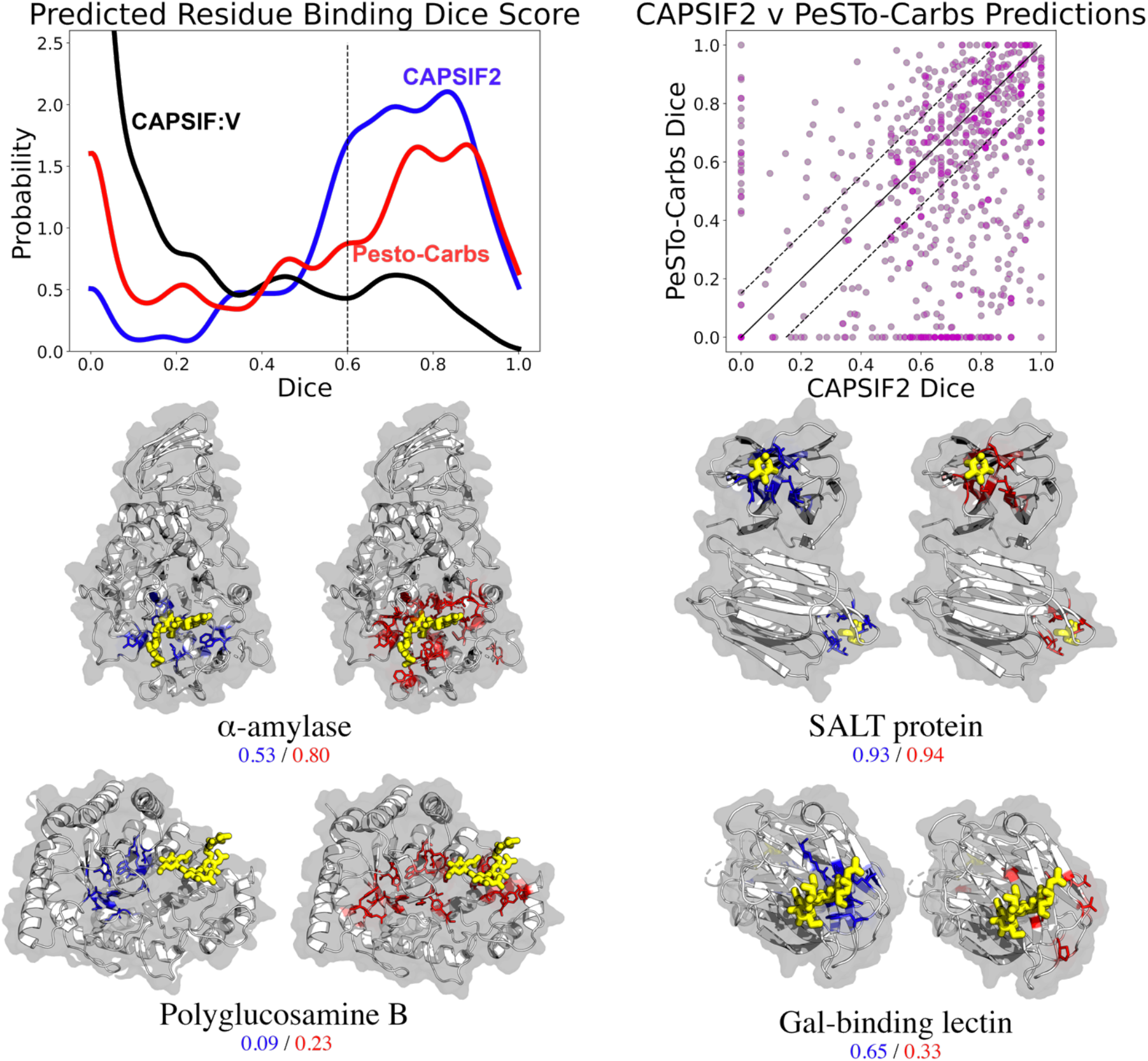
Comparison of CAPSIF2 and PeSTo-Carbs on residue-wise prediction tasks. (A) Distribution of Dice coefficient across prediction targets (proteins) for CAPSIF2 (blue), PeSTo-Carbs (red), and CAPSIF:V (black) on the DR test set. Densities smoothed with a Gaussian kernel density estimate (KDE, bandwidth *h =* 0.04) . (B) Per-target comparison of CAPSIF2 to PeSTo-Carbs. (C) Side-by-side comparison of carbohydrate (yellow) bound proteins (gray) predictions by CAPSIF2 (blue, left) and PeSTo-Carbs (orange, right) on α-amylase (1BAG), SALT protein (5GVY), polyglucosamine subunit B (4P7R), and galactose binding lectin (5XFD). Per-target Dice coefficients shown below.

We further show the results of specific targets in Figure 1C. In most of these cases, PesTo-Carbs and CAPSIF2 can successfully find the binding region, with varying accuracy; however, they both appear to fail on some targets, such as polyglucosamine B. This target notably has an observable pocket in the center of the structure, which CAPSIF2 and PesTo-Carbs incorrectly identifies as the binding region, wherein the experimentally solved oligosaccharide is proximal to the pocket.

### PiCAP accurately predicts carbohydrate binding and non-binding on experimental structures

Leveraging the same foundational network structure as CAPSIF2, we constructed the equivariant graph neural network (EGNN) named Protein interaction of Carbohydrate Predictor (PiCAP) with five additional layers to yield a single value prediction of whether a protein does or does not bind a carbohydrate. PiCAP assesses the spatial relationship of residues over an increasing context window, pooling the sequence into a fixed size 2D image, and providing a singular classification prediction based on that 2D representation. To our knowledge, PiCAP is the first DL model to assess protein-noncovalent binding of carbohydrates at a protein level.

We tested PiCAP on a holdout set based on sequence similarity, finding that PiCAP achieves an 89.6% balanced accuracy (BACC), with a 96.3% true positive rate (TPR) and 82.8% true negative rate (TNR) (Table 3). The ability to separate out carbohydrate binding (blue) and non-binding (red) proteins is further demonstrated in 2D t-distributed stochastic neighbor embedding (T-SNE) plots (Figure 2)^29^. Despite our best efforts, we do expect that the nonbinder dataset is likely contaminated with some carbohydrate binding proteins, therefore we must further discriminate PiCAP’s ability to predict on specific test set subsets.

**Table 3:** Metrics for PiCAP on the NoCAP test set and associated subsets with the number of proteins in parentheses. BACC is balanced accuracy. TPR is True Positive Rate TPR = TP / (TP + FP). TNR is True Negative Rate TNR = TN / (TN + FN).

| Test Set | Accuracy |
| --- | --- |
| NoCAP BACC (4,411) | 0.896 |
| NoCAP TPR (2,374) | 0.963 |
| NoCAP TNR (2,037) | 0.828 |
| TNR Ribosome (7) | 1.0 |
| TNR Holdout (92) | 0.902 |
| TPR ProGen Lysozymes (69) | 0.841 |
| TNR Designed Nonbinders (186) | 0.608 |
| Antibody BACC (50) | 0.562 |

**Figure 2:**
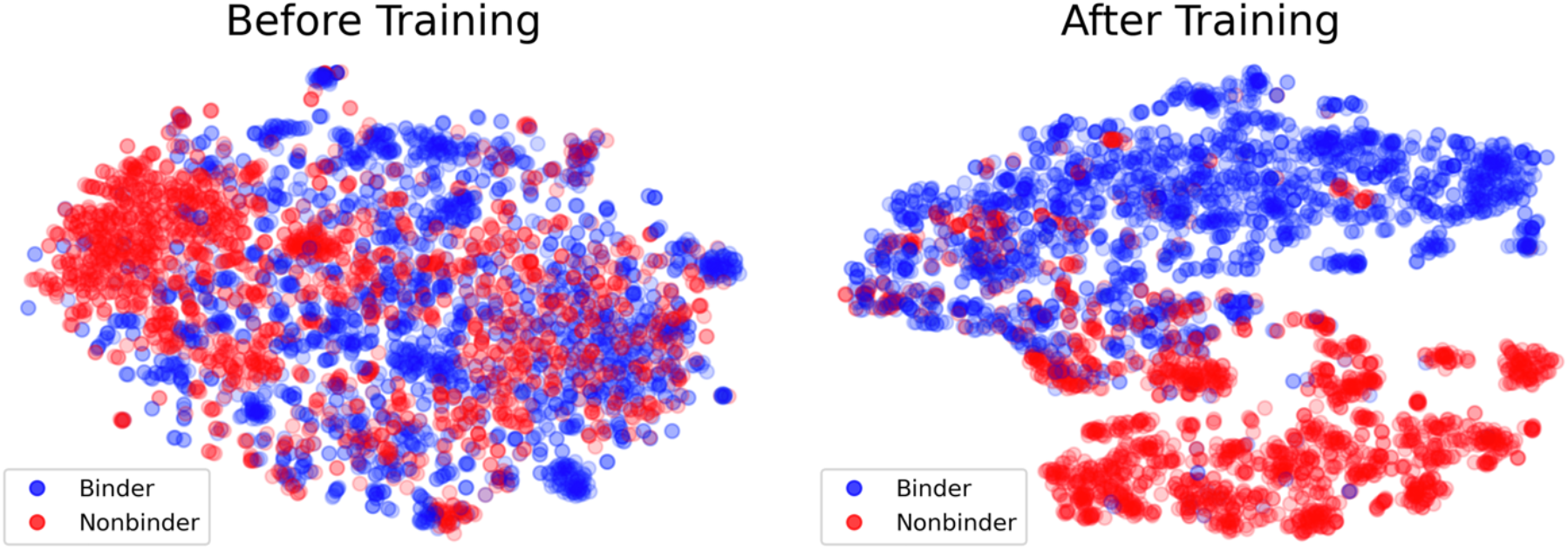
T-distributed stochastic neighbor embedding (T-SNE) diagrams of the PiCAP final layer embeddings of the NoCAP test set. (A) The randomly initialized model’s final layer output. (B) The final trained model’s final layer output.

When inspecting subsets of NoCAP (Table 3), we find PiCAP correctly predicts all the protein chains of the ribosome assembly as non-binders. We further have a holdout set of multiple proteins from various protein families, consisting of fatty acyl synthases, actin, myosin, and flippases, where PiCAP achieves an encouraging 90.2% accuracy on this negative subset. We observed that PiCAP performed well on designed lysozymes from the ProGen language model^24^ with an 84.1% accuracy. Contrarily, PiCAP achieves poor accuracy on computationally designed non-binder proteins, these being poor designs regarded as non-binding to the carbohydrate on the designed pocket, with an accuracy of only 60.8%. As a final test, we asked how our model performed on antibodies – specifically to identify antibodies that bind proteins or the glycans of glycoproteins. Of the 50 tested antibody structures, PiCAP achieved a 79% TNR and 33% TPR for a BACC of 56%. The antibodies and designed nonbinders are proteins hypervariably mutated at the binding site for specificity, which has the poorest performance of PiCAP, whereas PiCAP performs encouragingly on more evolutionarily and biologically defined proteins.

### PiCAP agrees with LectomeXplore and experimental evidence

Our NoCAP dataset for training and testing PiCAP comprises *experimentally* solved structures; therefore, we decided to investigate how our model performs on two datasets of *computationally predicted* structures. The first dataset is LectomeXplore published by Bonnardel et al., which identifies likely lectins across 37,794 organisms using a hidden Markov model (HMM) based on sequence and structural similarity.^22^ We also investigated the ganglioside interactome as published by Zhang et al., where they developed a high throughput assay to identify putative human proteins that interact with gangliosides.^23^ Both of these datasets have only sequence/UniProt gene IDs, therefore, for input into our algorithm, we used the predicted structures of the AF2 model proteomes^12^, only retaining confident segments of the structure (pLDDT larger than 70).

### LectomeXplore

The most closely related work to PiCAP is LectomeXplore, which identified putative lectins through sequence and structure homology. Unlike LectomeXplore, PiCAP does not limit proteins to be only of the lectin superfamily. We compared the likelihood of all predicted LectomeXplore lectins (greater than 0.25 confidence) present in the AF2 reference proteomes of two model species, *M. Musculus* and *H. Sapiens*, finding the agreement between the available AF2 structures of LectomeXplore and PiCAP to be 100% (225 of 225) for *M. Musculus* and 99.6% (229 of 230) for *H. Sapiens*. These results suggest a strong true positive rate (TPR) of PiCAP on the simplest class of sugar binding proteins.

### Ganglioside Interactome

Zhang et al. developed a high throughput method to identify proteins that interact with gangliosides. They created ganglioside probes with photoaffinity tags that covalently linked the probe to nearby proteins, and then they used mass spectroscopy and statistical methods to identify those proteins. They used six different probes in two different cell lines (A431 and SH-SY5Y), and for a total of nine experiments; we filtered the putative proteins by experiment. We selected the top 250 proteins above background from each experiment and removed CRAPome proteins.^30^ This identified 873 unique proteins across all nine experiments.^23^. As a high throughput method, and the first and largest of its kind, the error rates of their method have yet to be explored and cross-validated across other experimental methods. We therefore will use PiCAP to investigate the putative proteins of the ganglioside interactome work.

Of the 873 identified candidate ganglioside binding proteins, we were able to identify 848 proteins in the AF2 reference human proteome. PiCAP predicts 506 (60%) of these proteins as carbohydrate binders. Further, PiCAP also predicts that 988 of 3,500 putative non-ganglioside binding proteins (28%) as likely carbohydrate binders. Although these numbers at first suggest a substantial disagreement between our works, we see a strong positive increase in the fraction of proteins predicted as carbohydrate binders compared to the number of experiments that identified a binding protein (Figure 3A).

**Figure 3:**
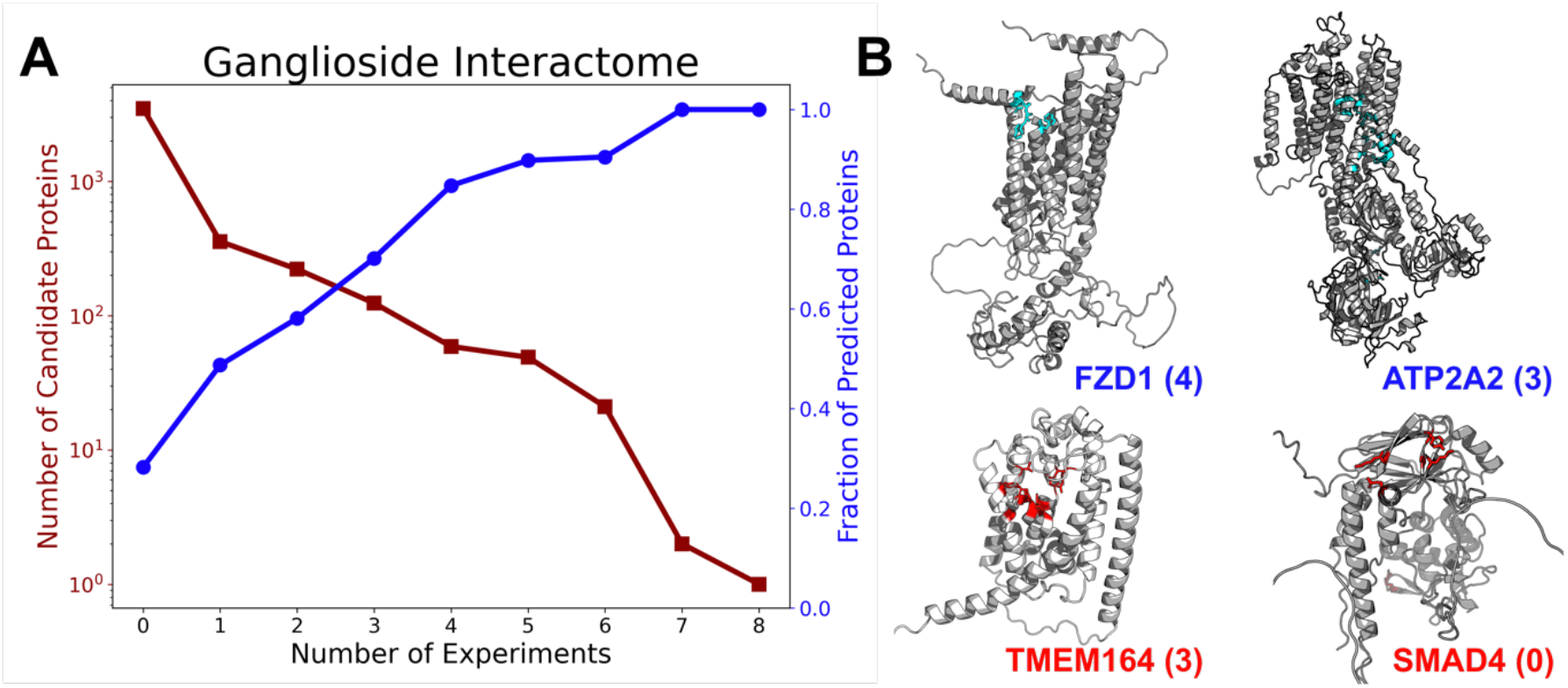
PiCAP validation against computational datasets. (A) Plot of Zhang et al. identified proteins across nine experiments alongside the fraction of proteins in each bin predicted as a carbohydrate binder by PiCAP. (B) PiCAP and CAPSIF2 predictions of selected ganglioside interactome proteins. The top row (blue) indicates proteins predicted as carbohydrate binders by PiCAP and bottom row (red) as proteins predicted as non-binders by PiCAP with the number of experiments the protein was identified by in parentheses. Highlighted residues in cyan (top column) and red (bottom column) are the predicted binding regions by CAPSIF2.

To explore the agreement and disagreement between our experiments, we selected four representative proteins: Frizzled-1 (FZD1), ATPase sarcoplasmic/endoplasmic reticulum Ca2+ transporting 2 (ATP2A2), transmembrane protein 164 (TMEM164), and mothers against decapentaplegic homolog 4 (SMAD4). FZD1 is involved in the Wnt signalling pathway and was identified by four of Zhang et al.’s experiments; PiCAP predicts FZD1 as a carbohydrate binder, and FZD1 was a subject of close scrutiny in the ganglioside interactome work^23^. ATP2A2 is an intracellular calcium/ATP pump and was identified by three experiments, and predicted as a carbohydrate binding protein by PiCAP. ATP2A2 has a specific role in ATP-mediated transport of calcium ions and likely little specific affinity for carbohydrates, let alone gangliosides^31^. TMEM164 is a regulator of ferropoptosis (iron mediated cell death) by an enzyme reaction with polyunsaturated fatty acids (PUFA) and was identified by three experiments^32^. PiCAP disagrees with the experimental results and predicts TMEM164 as a non-binder, which could indicate a potential error in the experimental evidence. Finally, SMAD4 is a transcription factor^33^, which was identified to not interact with gangliosides in all experiments, where PiCAP agrees and predicts the protein as a carbohydrate non-binder.

### PiCAP and CAPSIF2 can predict putative proteome scale interactomes

With PiCAP validated to an acceptable level, we sought to understand the protein-carbohydrate interactome with greater breadth than studied before. We chose three model organisms from the AF2 proteome datasets^12^, *E. coli, M. musculus*, and *H. sapiens*. Of the 4,363 proteins in the AF2 *E*.*coli* proteome, PiCAP yielded predictions on 4,339 accessible proteins and predicted 1,677 (39%) proteins as carbohydrate binders. Of the 21,615 proteins in the AF2 *M. musculus* proteome, PiCAP yielded predictions on 21,304 proteins and predicts 8,177 (38%) proteins as carbohydrate binders. Of the 20,650 proteins in the AF2 *H. sapiens* proteome, PiCAP yielded predictions on 20,067 proteins and predicts 7,029 (35%) proteins as carbohydrate binders (Figure 4A). We further provide the results of three additional model species: *Drosophila, C. elegans*, and *S. cerevisiae* in the supplemental information, without detailed analysis. We further analyze proteins of unknown function from *E. Coli* in Figure S1.

**Figure 4:**
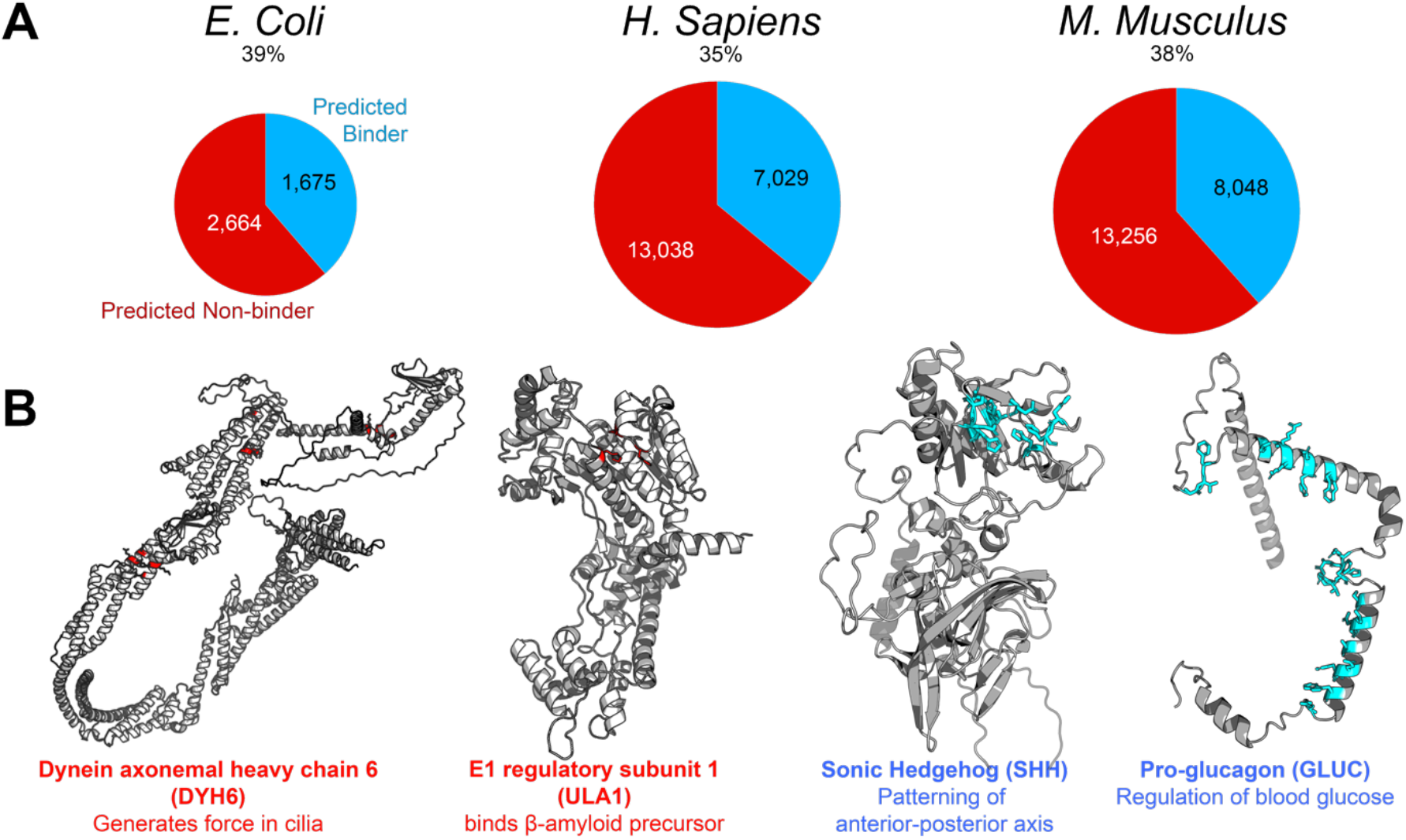
PiCAP predictions of proteomes. (Top) comparison of the fraction of proteins predicted as carbohydrate binders by PiCAP across three proteomes. (Bottom) PiCAP and CAPSIF2 predictions of four selected human proteins.

### Human Proteome

The primary proteome we analyzed was the AF2 human proteome (UP000005640), which contains 20,650 unique proteins with substantial resolution. PiCAP predicted 7,029, or 34%, of proteins to bind to carbohydrates. For comparison, the total number of lectins identified by LectomeXplore is 230, or 1.1% of the UniProt reference proteome^22^, and the number known by CaZy is 349, or 1.7%^21^. In contrast, the number of proteins experimentally identified as likely to bind gangliosides, a unique glycan family, is 873, or 4.2%^23^. To reconcile the differences between our work and the work of many others, we spot checked PiCAP’s predictions with several randomly selected proteins. We selected four proteins that appear representative of the distribution (Fig 4B).

PiCAP predicts sonic hedgehog (SHH), a protein implicated in cellular differentiation that can exist in the extracellular space to mediate the anterior-posterior axis, as a carbohydrate binding protein^34^. PiCAP additionally predicts pro-glucagon (GLUC) as a carbohydrate binding protein.

GLUC is a protein that becomes proteolyzed into constituent peptides, such as glucagon-like peptide 1 (GLP-1), which stimulate glucose-dependent insulin release, which can be mimicked by GLP-1 agonist pharmaceuticals such as dulaglutide (Trulicity) and semaglutide (Ozempic)^35^. PiCAP predicts E1 regulatory subunit 1 (ULA1), a protein that binds β-amyloid precursor protein, as a non-binder^36^. PiCAP additionally predicts dynein axonemal protein (DYH6), a protein implicated in microtubule-associated motor proteins, as a carbohydrate non-binder^37^.

## Discussion

We have demonstrated (1) an updated protein carbohydrate site identifier CAPSIF2 that outcompetes all current models on a generalized dataset and (2) a novel model named PiCAP that predicts *whether* a protein binds to carbohydrates or not. We validate our models against other models and datasets and applied to proteome scale analysis to garner more information about the protein-carbohydrate interactome.

CAPSIF2 boasts modest improvements in prediction accuracy on the original CAPSIF/TS90 dataset compared to CAPSIF:G and CAPSIF:V, but it underperforms PesTo-Carbs. CAPSIF2 however excels the most at a larger dataset containing ∼1k structures with substantially larger sequence variability, outcompeting all tested models. CAPSIF2 leverages a graph neural network operating on residues, using the same foundational approach as CAPSIF:G, while CAPSIF:V used a 3D voxelized CNN approach. PesTo-Carbs also leverages a graph neural network approach; however, it operates at an atom-wise level and only pools to the residue level late in the architecture. These graph architectures however have a similar level of parameters, where CAPSIF:G has 236K parameters, CAPSIF2 has 1.6M parameters, and PesTo-Carbs has 1.1M parameters; while CAPSIF:V has substantially more with 102M parameters.

We believe that the differences in performance are primarily not attributable to the architectures themselves, but rather the datasets. All models perform in a Matthews correlation coefficient (MCC) range from 0.55 to 0.63; thus, we attribute the largest differences to the stochastic training of these models and the slight variations in architectures. Structural protein-carbohydrate datasets are limited currently by the size of the PDB, as these interactions must be strong and stable to observe with experimental methods, where in physiology these interactions are often guided by avidity over affinity and/or enzymatic activity on the carbohydrates themselves. We believe larger datasets is only one part to improving these models, but improving the datasets with manual interrogation of all structures and with the identification of continuous biophysical pockets is necessary to improve the models’ performance.

To improve carbohydrate-protein structural datasets and improve our general biological understanding of the carbohydrate-protein interactome, we created PiCAP. PiCAP is the first model of its kind as it predicts carbohydrate binding of proteins independent of family/function – whether it be a cell surface protein for adhesion and communication or for metabolic enzymatics. The dataset we used to train PiCAP primarily separates known carbohydrate binders and proteins that are unlikely to bind to carbohydrates physiologically inside the cell – ranging from small molecule binders to cytoskeletal components. Although this approach is imperfect, it is the first attempt of this kind and leverages biophysical intuition of the cellular systems. Ultimately PiCAP achieves 89.6% accuracy on the experimental dataset with 1.8M parameters. PiCAP predicts most subsets of the test set with equivalent accuracy (designed lysozymes, cytoskeletal proteins, flippases, and fatty acyl binding proteins); however, it proves notably worse on designed-non binders and antibodies. The designed non-binders were created using Rosetta, where the binding pocket itself was designed but the remainder of the protein remained untouched^25^. These designs were labeled as non-binders by positive Rosetta binding energy scores – and never experimentally expressed nor tested. In a similar vein, to bind carbohydrates, antibodies use their hypervariable regions which are local regions that undergo somatic hypermutation. Our input to the protein is ESM2 embeddings, which uses full sequence context to extract a large 1280-dimensional embedding of each residue. As the ESM2 model is only trained and tested on biological proteins, the signal specificity of the binding pocket sequence of the designed non-binders may be masked by its more evolutionarily conserved residue, leading PiCAP to predict these non-biological proteins as carbohydrate binders. PiCAP studies protein sequence and structural information together, indicating PiCAP as a strong candidate for proteome wide studies of protein-carbohydrate interactions.

Since PiCAP performed well on NoCAP data, we sought to validate the model against other methods that predicted carbohydrate binders: the ganglioside interactome and LectomeXplore. While we saw only 60% of proteins in the ganglioside interactome as positive, after closely evaluating a subset of the data, we reconciled the difference with the error of the high throughput experimental method. Although PiCAP appears to disagree with a good fraction of the ganglioside dataset, it has a strong linear relationship with the high throughput experiments. The more experiments that identified a protein, the higher likelihood that PiCAP predicted the protein as a carbohydrate binder. We further observe strong agreement between LectomeXplore and PiCAP, with an average of 95% agreement across three model species.

With experimental and computational validation, we then leveraged PiCAP against the AF2 proteome datasets. PiCAP predicts 35∼40% of all proteins in three biological model species to be carbohydrate binding proteins – the highest prediction to date. As carbohydrates are ubiquitous across all species and are the foundational building block of energy storage and integral to most all extracellular communication, it is unsurprising for such a high fraction of proteins bind to carbohydrates. PiCAP results can be further validated by proteomic evaluation by experiments such as the pull-downs from Zhang et al.^23^ or liquid glycan arrays^38,39^. The computational predictions can help elucidate more functionality of proteins and provide a larger context to their roles inside the cell and the suggestion of more protein moonlighting than previously understood. Despite their biophysical importance in most all cellular functions, carbohydrates remain elusive with few studies determining the exact extent of protein-carbohydrate interactions. Our work expands to all proteins/carbohydrates in an agnostic manner that abstains from any limits on protein family or carbohydrate species. We released the results of CAPSIF2 and PiCAP of six model system proteomes for all proteins for open-source scientific use. Additional steps can now be taken for the ultimate goals to design proteins to carbohydrate and glycoprotein targets for therapeutic purposes. Firstly, we encourage the expansion of this work or LectinOracle^40^ or GlyNet^41^ to predict carbohydrate species to all carbohydrate binding proteins. One simple step would be to predict whether proteins bind to just a specific species of carbohydrate – such as chitins or sialic acids. Another step would a high throughput computational docking of those carbohydrate species to the identified proteins, using CAPSIF2 or PesTo-Carbs^17^ or DeepGlycanSite^18^ to identify an initial hypothesis to feed GlycanDock^42^, or directly *de novo* with programs like DiffDock^13^, RosettaFold-All Atom (RF-AA)^43^, AlphaFold3^11^, or Boltz-1^44^ (although there are currently no validation studies testing whether these methods provide high accuracy on carbohydrate-specific docking).

In addition, all these methods leverage deep learning techniques. Deep learning methods require multitudes of data, and although we were able to demonstrate impressive results on low accuracy/messy data, we believe a clean dataset is integral and necessary for the future of this field. A better annotated set of proteins that do not bind carbohydrates would be helpful, as well as all structural proteins to have all ligands together, where currently there is a high redundancy in protein structures with slightly different ligands or crystallization techniques, which reduce the accuracy of the test metrics in comparing CAPSIF2 and PesTo-Carbs. We also believe tandem experiments, such as those done by Zhang et al.^23^ or selective exo-enzymatic labeling (SEEL) glyco-engineering high throughput methods^45^ to validate these models could further demonstrate a larger wealth of carbohydrate binding proteins, alongside their specificity, allowing for further annotation of the genome on a large scale.

## Methods

### Dataset

Carbohydrate-binding proteins were selected by combining multiple datasets. We selected carbohydrate binding antibodies from SAbDab,^26^ all experimentally solved proteins from UniLectin^19^ (with and without bound carbohydrates), the CAPSIF dataset,^16^ and most notably, the DIONYSUS dataset.^20^ Further, we included the computationally designed and experimentally viable lysozymes from ProGen,^24^ with structures predicted by the Colab distribution of AlphaFold2.^46^

There are several datasets of protein-carbohydrate interactions; however, there is no dataset of proteins that do not bind to carbohydrates, so we constructed one (Table 1). In the creation of such dataset, an intrinsic difficulty is that it is not possible to prove that a protein does not bind to a carbohydrate of any kind; therefore, we selected proteins that biophysically have low likelihood to bind to carbohydrates due to their function or location inside the cell. The experimentally solved proteins selected were primarily chosen as small molecule binding proteins, DNA binding proteins, nuclear pore complex proteins, serine proteases, cytoskeletal proteins, aminotransferases, flippases, fatty acid binding proteins, selected antibodies (antibodies), and ribosomal proteins. In addition to these proteins, Luo et al. computationally constructed a dataset of carbohydrate non-binder proteins with the Rosetta software^25^.

Small molecule data constitutes the largest portion of the non-binders (∼18k pdbs), as we used the PDB-Bind 2020 dataset.^27^ Some proteins in the PDB-Bind dataset contain carbohydrates as the ligand, in which case we identified those ligands using PyRosetta^47^ and removed them from the non-binder dataset and added them to the binder dataset. Antibodies were selected using the SAbDab dataset by finding all proteins that were bound to proteins or nucleic acids and further filtering to structures not containing any carbohydrates in the structure nor an NX(S/T) motif in the antigen.^26^ Ribosomal proteins were selected from the bacterial ribosome structure.^48^ The remainder of protein structures were selected by inspection from the RCSB PDB.^49^

After combining the datasets and adjusting for duplicate PDBs across different datasets, the final NoCAP dataset contains 30,849 total unique protein structures. Of these structures, 9,608 bind to carbohydrates, with 6,724 having an experimentally bound carbohydrate. Of the 21,412 non carbohydrate binders, 17,191 have an experimentally resolved small molecule bound to it, leaving 4,221 as nonbinders. To encourage generalizability to minor errors in structure predictions, we also reconstructed the 12,021 shortest sequence proteins of the 30,849 with the Colab implementation of AlphaFold 2^46^, where we only kept the 11,042 of predicted structures with a pLDDT greater than 80.

### Preprocessing

With our dataset, we desired to leverage both sequence and structural information to predict carbohydrate-binding capabilities of proteins. Family information of sequence similarity can strongly indicate carbohydrate binding capabilities, while structural motifs can be present across protein families for carbohydrate binding, and we desire our method to identify both. We extracted the sequence and the Cβ positions of all protein residues (Cα for glycine) using PyRosetta.^47^ Next, we used ESM2^15^ to provide a high-dimensional sequence embedding for each protein residue of each protein chain. We labeled protein residues that were within 4.2 Å of a non-covalently bound carbohydrate (or small molecule) as a binding residue.

Most previous work has used single protein chains for protein-carbohydrate predictions^16,17^; however, many proteins only exist in the context of multiple chains. For this reason, we preprocessed all protein structures with all chains in the PDB file, except the initial CAPSIF dataset and antibodies. To limit the redundancy of the training set, we used MMseqs to cluster protein sequences by 60% sequence identity into distinct clusters for training/testing.^50^ We then split the clusters into an 80/5/15 train/validation/test, maintaining the same proteins from CAPSIF remain in the same dataset distribution. This left 24,957 structures in training, 1,479 structures for validation, and 4,413 structures for testing.

### Secondary validation set

We have a primary dataset of carbohydrate-protein binding; however, we need to demonstrate the ability of PiCAP to predict outside of the crystally solved structures. To do this, we gathered all UniProt^51^ accession codes from Zhang et al.^23^ and LectomeXplore^22^ and matched them to the AF2 publicly accessible organism proteomes. This captured 848 of 878 (97%) of putative ganglioside binding proteins and 3,400 of 4,335 (78%) of non-ganglioside binding proteins. LectomeXplore uses sequence and structural protein information, alongside infectious pathogens that affect these species, and lists all reference sequences and structures (UniProt, ensembl, NCBI, RCSB, etc.) with severe redundancy. Therefore, for a direct quantitative comparison, we therefore used only those that existed singly as UniProt values inside the reference proteomes of AF2. We used the confidence metric of 0.25 for identification of lectins, which yielded 230 human proteins and 225 mouse proteins.

For our proteome analysis, we used the AF2 publicly accessible organism proteomes.^12^ AF2 generates structures with an internal confidence metric called pLDDT, where low confidence regions will have pLDDTs under 70. We therefore performed analysis and studies on AF2 protein regions with high confidence, or residues with greater than 70 pLDDT, independent of structural continuity. We applied the analysis to the following model organisms: *E. Coli, M. Musculus*, and *H. Sapiens*. We further provide the results of the full-length sequence, independent of pLDDT in Supplemental information alongside the results of three other model organisms: *C. elegans, D. melanogaster*, and *S. cerevesia*.

### Architectures

We fed the residue coordinates and sequence embeddings into the CAPSIF2 and PiCAP architectures (Figure 1A) into the main model block, which uses a message passing equivariant graph neural network (EGNN) of equivariant graph convolutional layers (EGCL).^52^ Each layer sums the outputs of a multilayer perceptron (MLP) that inputs the features of the central node and the features of all of its neighboring nodes and the edge attributes of the neighbors. Following Ingraham et al.^53^, the edge attributes are a radial basis function (RBF) of the distance, the orientation, and direction of the neighboring residues.

CAPSIF2, a carbohydrate binding residue predictor, has 12 residual ECGLs with an embedding dimension of 128 (Figure 5). The neighborhood context window is fixed at the 16 nearest neighbors. After the graph convolutions, each residue is passed to a two-layer dense decoder, finally outputting the carbohydrate-binding likelihood of each residue. CAPSIF2 contains 1,600,387 parameters.

**Figure 5:**
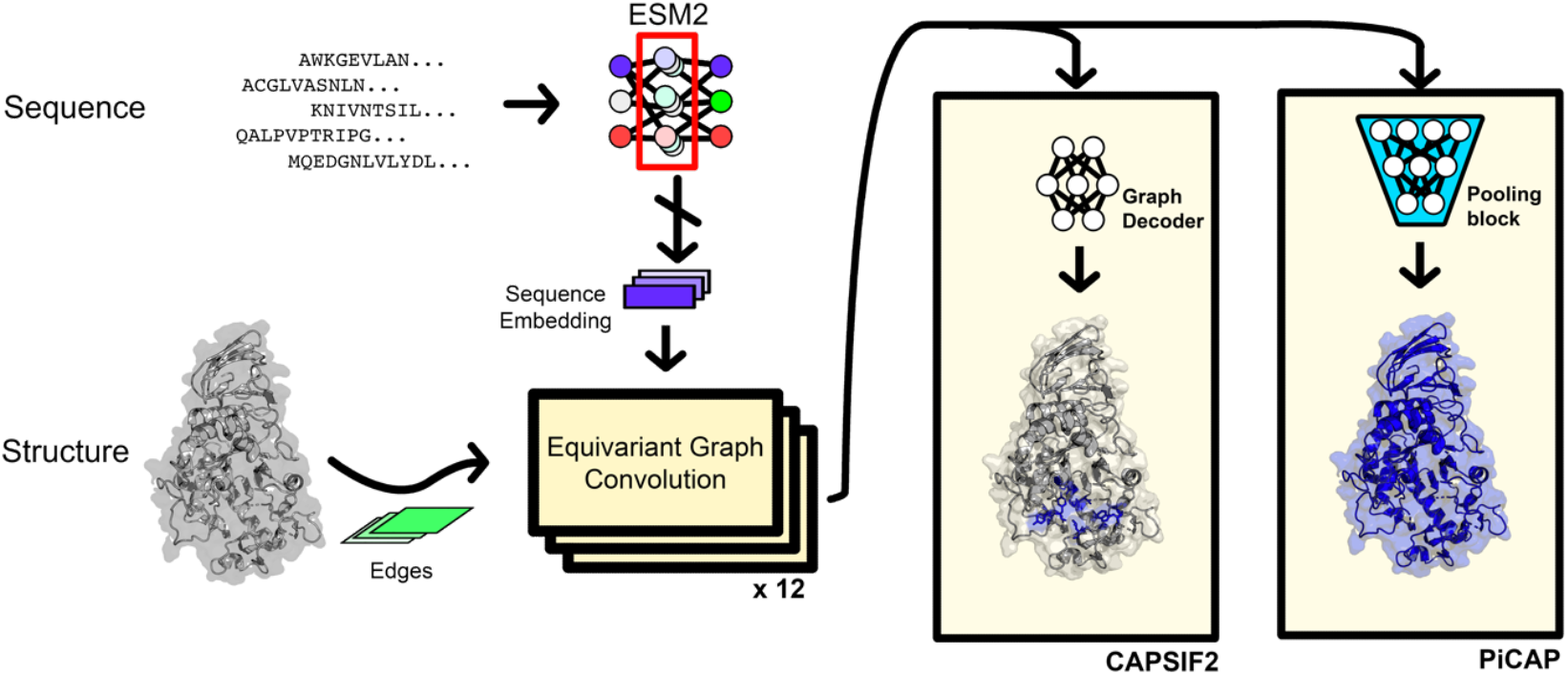
Architectures of CArbohydrate Protein Site IdentiFier 2 (CAPSIF2) and Protein interaction of CArbohydrate Predictor (PiCAP).

PiCAP, a predictor of whether a protein binds to carbohydrates, has 12 total residual EGCLs with an embedding dimension of 128 and leverages an increasing neighborhood context window for information propagation, as inspired by PeSTo^54^ and PeSTo-Carbs^17^. The first three layers use the 10 nearest neighbors, layers 4 to 6 use the 20 nearest neighbors, layers 7 to 9 use 40 neighbors, and layers 10 to 12 use 60 neighbors. (Figure 5). The model specific block is a pooling block that uses an adaptive pool to truncate or slightly expand the size of the protein to a fixed length (150), where the model then uses two convolutional layers and three dense layers to predict the likelihood of a protein to bind to carbohydrates. PiCAP contains 1,798,895 parameters.

### Training

We trained both models using two cycles: small molecule binding residue prediction and the model specific task (protein or residue level predictions). The first training cycle used the CAPSIF2 base architecture with randomized initial weights ∼*N*(0,0.02) for the residue level prediction. The model was trained for a maximum of 1,000 epochs, with training prematurely stopped once the validation loss did not decrease after 35 epochs. This training cycle had a learning rate of 2 × 10^−6^ and a weight decay of 10^−7^ with the Adam optimizer with the loss function *L* = 1 − *d*, where *d* is the Dice-Sorenson coefficient (also known as the F1 score) and a batch size of 1. To improve model generalization, each epoch sampled a single protein from every training cluster available from the small molecule dataset. The smallest 12,000 protein sequences were modeled structurally with the colab distribution of AF2,^10,46^ and if the selected protein was available via AF2, we selected the crystal structure 40% of the time and the AF2 structure 60% of the time.

For the second training cycle, CAPSIF2 used the same architecture as the first training cycle and required no randomization. CAPSIF2 was trained only on proteins with experimentally determined carbohydrate binding sites with learning rate of 2 × 10^−5^ and weight decay of 10^−6^ with the Adam optimizer and the same loss function of *L* = 1 − *d*. Similar to the first training cycle, we randomly selected an available AF2 structure 60% of the time.

For the second training cycle, PiCAP used the weights where available from the first training iteration of CAPSIF2 and randomized weights for the model specific block ∼*N*(0,0.02). PiCAP was trained on the entire training set for binary classification with a learning rate of 2 × 10^−5^ and weight decay of 10^−6^ with the Adam optimizer and binary cross entropy (BCE) loss function. Similar to the first training cycle, we randomly used an available AF2 structure 60% of the time.

## Data Availability

Data, code, and datasets are available at Github, where CAPSIF2 and PiCAP can be run at: https://github.com/Graylab/picap. We further provide a webserver on ROSIE where CAPSIF2 and PiCAP can additionally be run: https://r2.graylab.jhu.edu/apps/index.

## Author Contributions

S.W.C. conceived the project, performed the research, analyzed data, wrote the manuscript, and created all figures. R.L.S. supervised the project, analyzed data, and wrote the text. J.J.G. conceptualized and supervised the project, analyzed data, and wrote the text.

Funding

This work was supported by NIH R35-GM141881 (JJG and SWC), and NIH R01-AI162381 (JJG and SWC).

## Acknowledgements

We thank Yijie Luo and Fabio Parmeggiani (Univeristy of Bristol) for the non-binder dataset. All model training and testing was performed on Johns Hopkins Advanced Research Computing at Hopkins (ARCH). We thank Sergey Lyskov for assisting in the implementation of the CAPSIF2 and PiCAP web server on ROSIE.

## Notes

https://github.com/Graylab/picap

### Competing Interest Statement

The authors have declared no competing interest.

### Summary of Updates

Change of title Small typos in main and supp

