## Supplementary File 1 for "Predictions from Deep Learning Propose Substantial Protein-Carbohydrate Interplay"

### Supporting Information

#### The Proteome as a Lectome: Predictions from Deep Learning Propose Substantial Protein-Carbohydrate Interplay

Samuel W. Canner[1], Ronald L. Schnaar[2,3], Jeffrey J. Gray[1,4]
[1]Program in Molecular Biophysics, Johns Hopkins University, Baltimore, MD, United States
[2]Department of Pharmacology and Molecular Sciences, Johns Hopkins University School of Medicine, Baltimore, Maryland, United States
[3]Department of Neuroscience, Johns Hopkins University School of Medicine, Baltimore, Maryland, United States
[4]Department of Chemical and Biomolecular Engineering, Johns Hopkins University, Baltimore, MD, United States

### Dataset Description

We provide Supplemental File 1 as an Excel document (xlsx). This Excel document contains all prediction information of PiCAP and CAPSIF2 on the AlphaFold 2 proteomic data[1]. Since PiCAP can produce false negative binders, the separate predictions of CAPSIF2 in all cases may assist in hypotheses of known carbohydrate and small molecule binding proteins. In addition, we show predictions on all proteins in the dataset, even in cases with low pLDDT, although our analysis in the primary text only analyzes predictions of proteins with greater than an average of 70 pLDDT. In all sheets, we provide the following columns:

- UniProt Entry
- Common Gene Name (Entry_Name)
- Protein_name
- Gene Ontology terms
- PiCAP prediction on only residues with greater than 70 pLDDT
- CAPSIF2 predicted binding residues on residues with greater than 70 pLDDT

### Model Hyperparameterization

To optimize performance on a neural network, we assessed multiple hyperparameters in both models to achieve their performance. We focused primarily on the following hyperparameters: embedding dimension, *k*-nearest neighbors (knn), and number of layers. For simplicity, we treat each network as a series of four (4) blocks, composed of a certain number of layers where we vary knn per block.

#### CAPSIF2 parameterization

In our previous work on CAPSIF:G, we used one-hot encodings of amino acid type and biophysical properties with simple edge embeddings. To contain more information, in this work, we altered the node features to ESM2 embeddings and edges. With these input features, we then focused on the size and depths of the network.[2] A full account of all tested hyperparameters is listed below in Table S1. We selected CAPSIF2 as the model that performed the best on the DR test set, which was composed of 12 layers with a static number of k nearest neighbors of 16.

**Table S1: Performance of various CAPSIF2 models on the Dionysus Residue (DR) test set.** Dice and Matthews correlation coefficient (MCC) are as defined in the main text. Boldface indicates the best performance in each metric. Selected CAPSIF2 model is highlighted in yellow.

| Layers per block | KNN per block | DR Dice | DR MCC | TS90 Dice |
|---|---|---|---|---|
| 3 | 6,6,6,6 | 0.541 | 0.542 | 0.533 |
| 3 | 8,8,8,8 | 0.477 | 0.484 | 0.431 |
| 3 | 8,12,16,20 | 0.391 | 0.407 | 0.366 |
| 3 | 6,10,14,18 | 0.528 | 0.528 | 0.572 |
| 3 | 10,20,40,60 | 0.289 | 0.312 | 0.364 |
| 3 | 16,16,16,16 | **0.573** | **0.574** | 0.616 |
| 3 | 20,20,20,20 | 0.498 | 0.496 | 0.575 |
| 4 | 8,12,16,20 | 0.566 | 0.567 | **0.639** |
| 4 | 6,10,14,18 | 0.408 | 0.419 | 0.319 |
| 4 | 8,8,8,8 | 0.491 | 0.486 | 0.548 |
| CAPSIF:V | N/A | 0.226 | 0.202 | 0.608 |

#### PiCAP parameterization

We followed the same methodology as CAPSIF2 to identify the strongest performing PiCAP model parameters. Our hyperparameter search is provided below in Table S2. The decision on which model performed strongest was less straightforward than CAPSIF2, as all multiple models performed strongly across the NoCAP test set. We selected a model that performed well across most metrics placing just below the top of every other category to encourage generalizability, as some of the top performing models were prone to overfitting and unstable predictions. The chosen PiCAP model consisted of 12 layers, with the knn gradually increasing from 10 to 60 neighbors across the layers.

**Table S2: Performance of various PiCAP models on the NoCAP test set.** BACC is balanced accuracy. TPR is True Positive Rate TPR = TP / (TP + FP). TNR is True Negative Rate TNR = TN / (TN + FN).

| Layers per block | KNN per block | cutoff | NoCAP BACC | NoCAP TPR | NoCAP TNR | Nonbinders TNR | Ribosome TNR | Holdout TNR |
|---|---|---|---|---|---|---|---|---|
| 3 | 6,6,6,6 | 0.94 | 0.85 | 0.87 | 0.83 | 0.624 | **1.0** | 0.857 |
| 3 | **8,8,8,8** | **0.33** | 0.892 | 0.964 | 0.82 | 0.667 | 0.857 | **0.929** |
| 3 | 20,20,20,20 | 0.21 | 0.856 | 0.88 | 0.833 | 0.683 | **1.0** | 0.429 |
| 3 | 10,20,40,60 | 0.23 | 0.896 | 0.963 | 0.828 | 0.608 | **1.0** | 0.902 |
| 3 | 6,10,14,18 | 0.99 | 0.779 | 0.927 | 0.631 | 0.656 | 0.857 | 0.571 |
| 4 | 6,6,6,6 | 0.77 | 0.885 | **0.976** | 0.794 | 0.731 | **1.0** | 0.857 |
| 4 | 8,8,8,8 | 0.19 | **0.897** | 0.951 | **0.842** | 0.704 | **1.0** | 0.857 |
| 4 | 6,10,14,18 | 0.32 | 0.861 | 0.974 | 0.745 | 0.134 | 0.857 | 0.643 |
| 4 | 8,12,16,20 | 0.84 | 0.877 | 0.954 | 0.801 | **0.785** | **1.0** | 0.643 |

### Proteomic Data

In Supplemental File 1, we provide a list of all proteins from six organisms. Here we list the overall metrics of the three organisms in Supplemental File 1 that were not discussed in the main text. PiCAP predicts that in *C. elegans* (nematode worm) 9,278 of the 19,227 proteins (48%) bind carbohydrates. PiCAP predicts that in *D. melanogaster* (fruit fly) 5,248 of the 13,351 proteins (39%) bind carbohydrates. PiCAP predicts that in S. *cerevisiae* (yeast) 1,749 of the 5,849 proteins (29%) bind carbohydrates.

#### E. Coli Unknown Proteins

PiCAP is the first algorithm for the prediction of proteins that bind to carbohydrates; therefore, we used PiCAP to predict novel functions of proteins. We therefore used PiCAP to discern potential carbohydrate binding functionality in proteins of unknown function, according to gene ontology (GO) in the UniProt (as of Jan 29, 2025).[3] In *E. Coli*, there are 393 proteins of unknown functionality, and PiCAP predicts that 138 (35%) of these proteins as carbohydrate binding proteins. Three of the positive predicted binding proteins are shown in Figure S1: P28915 (YBFC), P45581 (STFP), and P36682 (YACH). The AF2 model of YBFC adopts a β-sandwich fold, which is a canonical fold of lectins, with CAPSIF2 predicted residues on the β-sandwich. The AF2 model of STFP appears to have a minimally ordered tail with an ordered β-sandwich domain, where CAPSIF2 predicts several residues to bind carbohydrates, which could indicate a membrane protein for binding to the extracellular space. The AF2 model of YACH has a primary domain of α-helices and β-strands with some floating alpha helices which may be part of a different domain; however, CAPSIF2 does not predict any residues to bind, and PiCAP only predicts a 72% carbohydrate-binding likelihood.

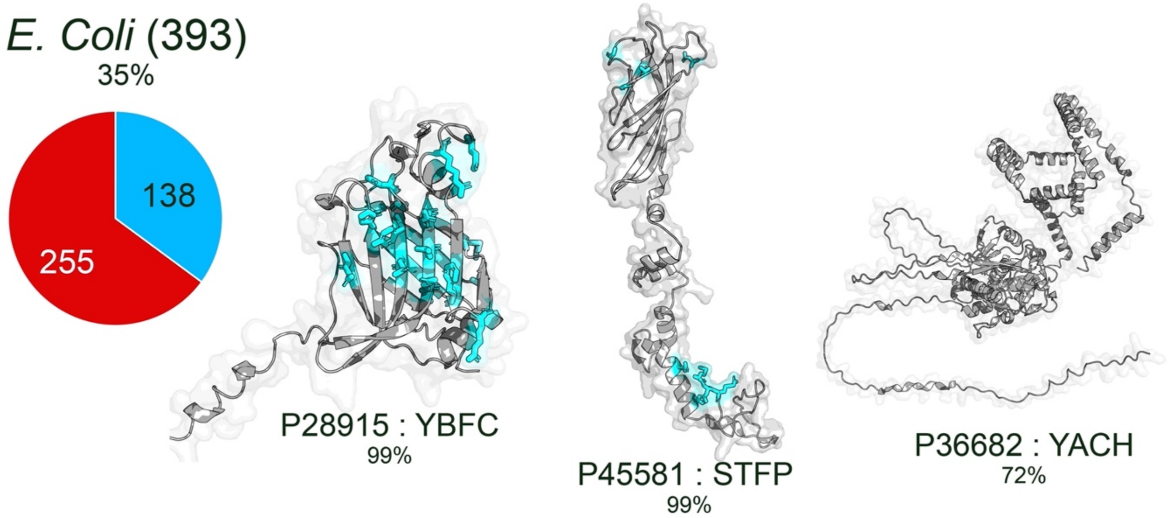

**Figure S1: PiCAP predictions of *E. Coli* unknown function proteins.** (A) comparison of the fraction of proteins predicted as carbohydrate binders by PiCAP across three proteomes. (B) PiCAP and CAPSIF2 predictions of three selected *E. Coli* proteins with PiCAP prediction provided.

1. Varadi, M. *et al.* AlphaFold Protein Structure Database in 2024: providing structure coverage for over 214 million protein sequences. *Nucleic Acids Res* 52, D368–D375 (2024).
2. Canner, S. W., Shanker, S. & Gray, J. J. Structure-based neural network protein–carbohydrate interaction predictions at the residue level. *Frontiers in Bioinformatics* 3, (2023).
3. Bateman, A. *et al.* UniProt: the Universal Protein Knowledgebase in 2023. *Nucleic Acids Res* 51, D523–D531 (2023).
